# Transcriptome-based genome-wide analysis reveals hybridization dynamics and genetic structure of Japanese giant salamanders

**DOI:** 10.64898/2026.05.26.727823

**Authors:** Takeshi Igawa, Sumio Okada, Mifuyu Sera, Rika Takagi, Masaki Yamazaki, Zenkichi Shimizu, Hidemasa Bono, Yoshihiro Omori

**Affiliations:** Graduate School of Integrated Sciences for Life, Hiroshima University, Higashi-Hiroshima, Japan; Amphibian Research Center, Hiroshima University, Higashi-Hiroshima, Japan; Seto Inland Sea Carbon-neutral Research Center, Hiroshima University, Higashi-Hiroshima, Japan; Hanzaki Research Institute, Asago, Japan; Nabari City Board of Education, Culture and Lifelong Learning Office; Mie Natural History Research Group, Matsuzaka, Japan; Genome Editing Innovation Center, Hiroshima University, Higashi-Hiroshima, Japan

**Author notes:** Corresponding authors Takeshi Igawa, Ph. D., Amphibian Research Center, Hiroshima University, Higashi-Hiroshima, Japan, Yoshihiro Omori, Ph. D., Laboratory of Functional Genomics, Graduate School of Integrated Sciences for Life, Hiroshima University, Higashi-Hiroshima, Japan.

## Abstract

The Japanese giant salamander (*Andrias japonicus*), an apex predator and a Special Natural Monument in Japan, is threatened by hybridization with introduced Chinese giant salamanders (*Andrias davidianus*). This hybridization has caused genetic introgression and expansion of hybrid populations, posing a serious conservation risk. Because morphological identification of hybrids is occasionally unreliable and current genetic methods rely on limited markers, a genome-wide approach is required. However, the extremely large genome (∼50 Gb) of giant salamanders has hindered whole-genome analyses.

In this study, we conducted transcriptome-based analyses of Japanese giant salamanders, Chinese giant salamanders, and their hybrids, generating RNA-seq data from 34 individuals. A total of over 419,000 SNP candidates were identified, from which 4,457 high-confidence SNPs in highly expressed genes were selected for analysis. Population structure analyses for Nabari colony revealed that hybrid individuals form two major groups, corresponding to different degrees of genetic contribution from Japanese and Chinese lineages. Most hybrids were inferred to be F2 or backcross individuals, while F1 hybrids were rare. Mitochondrial analysis indicated that all hybrids possessed Japanese-type mitochondrial genome, suggesting male-mediated introgression from Chinese salamanders.

Differential expression analysis revealed enhanced stress-response pathways in hybrids and stronger antiviral responses in Japanese individuals. Using the axolotl genome as a reference, we constructed a virtual chromosomal map, identifying large haplotype blocks and supporting recent hybridization with limited recombination. This study provides a genome-wide framework for understanding hybridization dynamics and supports future conservation and evolutionary studies.

## Introduction

The Japanese giant salamander (*Andrias japonicus*) is the largest amphibian species in Japan and acts as an apex predator, occupying the top trophic position in its river ecosystem (Duret et al. 2026). It is designated as a Special Natural Monument in Japan due to its biological distinctiveness, including its extreme body size (maximum total length of 150 cm) and longevity, as well as its threatened conservation status resulting from habitat loss and disease, and its cultural significance as an iconic national species (Matsui et al. 2008; Taguchi 2009; Une et al. 2012; Takaya et al. 2023). In Asia, three additional species of the genus *Andrias* are recognized as Chinese giant salamanders: *Andrias davidianus*, *Andrias sligoi*, and *Andrias jiangxiensis* (Turvey et al. 2019). These East Asian giant salamanders diverged relatively recently, approximately 8.3 million years ago, and are capable of hybridization (Turvey et al. 2019). Another member of the family Cryptobranchidae, the North American hellbender (*Cryptobranchus alleganiensis*), is distributed in North America. This species belongs to the genus *Cryptobranchus* and diverged from the genus *Andrias* approximately 60 million years ago (Turvey et al. 2019).

In China, the Chinese giant salamander is widely recognized as a distinctive food item with a unique texture (He et al. 2018), and it is supplied mainly through aquaculture. In addition, mucus secreted from its skin has been reported to be useful as a biomedical adhesive for postoperative wound closure, highlighting its potential for medical applications (Deng et al. 2019). On the other hand, wild populations of Chinese giant salamanders are also rare in China, and their conservation has been strongly advocated (Wang et al. 2004).

Spontaneous hybridization often occurs in amphibian species, and genome introgression has a significant impact on subsequent evolution (Komaki et al. 2015). On the other hand, hybridization caused by the artificial introduction of alien species or populations is also known to pose a major threat to environmental conservation. Hybridization between Japanese giant salamanders and introduced Chinese giant salamander (*A. davidianus*) in Japanese rivers has led to a rapid decline of the native and genetically pure Japanese giant salamander populations (Takahashi and Shimizu 2026). Moreover, hybrid individuals have expanded their distribution beyond the original habitats of the native species, creating an emerging environmental concern (Yoshikawa et al. 2012). Recent studies have revealed that Chinese giant salamanders imported into Japan for food consumption were released into Japanese rivers, such as the Kamo River in Kyoto, where they hybridized with native Japanese giant salamanders (Nishikawa et al. 2024). As a result, large numbers of hybrid individuals are now present in these river systems. Consequently, the genetic contribution of the Chinese lineage has been increasing at a faster rate within hybrid populations.

Hybridization patterns among closely related species are generally classified into six types (Allendorf et al. 2001). Among these, Type 6 describes a scenario in which hybridization between an introduced species and a native species leads to extensive genetic introgression, ultimately resulting in the extinction of the native species.

A well-known example of this pattern in Japan is the regional extinction of the native Japanese rosy bitterling (*Rhodeus ocellatus kurumeus*) following the invasion of the continental rosy bitterling (*Rhodeus ocellatus ocellatus*) (Kawamura et al. 2001; Kawamura 2005). In this case, despite the initially small population size of the introduced species, introgression of its genetic material into hybrid populations increased rapidly. This was attributed to the higher reproductive success and survival rate of the introduced species compared with the native species. As a result, the native rosy bitterling has become nearly extinct in Honshu and Shikoku.

On the other hand, ornamental goldfish, which are mainly derived from wild Chinese crucian carp, are capable of hybridizing with wild Japanese crucian carp. Indeed, genomic analyses have revealed that some goldfish varieties have a history of hybridization with Japanese crucian carp, raising concerns about their impact on environmental conservation (Chen et al. 2019; Kon et al. 2020). Another example is found in New Zealand, where the native Pacific black duck (*Anas superciliosa*) has been driven to functional extinction through hybridization with the introduced mallard (*Anas platyrhynchos*), and conservation efforts are now focused on managing hybrid populations (Allendorf et al. 2001). Careful attention is therefore required to prevent the native Japanese giant salamander from facing a similar fate.

Identification of hybrid individuals of giant salamanders has traditionally relied primarily on morphological characteristics. Chinese giant salamanders often exhibit a grayish body coloration, which differs slightly from the typical brown coloration with black mottling observed in Japanese giant salamanders. In addition, Chinese giant salamanders tend to have a flatter head shape, and the arrangement of small tubercles on the head differs somewhat from that of Japanese giant salamanders (Hara et al. 2023). However, these morphological traits vary considerably depending on habitat and individual variation, and therefore cannot serve as definitive diagnostic criteria. Moreover, in hybrid individuals, these characteristics often appear intermediate between the two species, making morphological identification even more difficult. Consequently, alternative methods for reliable identification are strongly needed. Genetic identification of giant salamanders is currently based on a limited number of polymorphic genetic markers (Yoshikawa et al. 2011; Yoshikawa et al. 2012). However, reliance on such a small set of markers carries a relatively high risk of misclassification. These limitations highlight the need to establish a genome-wide approach for detecting genetic polymorphisms. To date, genomic studies within the family Cryptobranchidae remain limited. Transcriptome analyses have been reported for the Chinese giant salamander (Geng et al. 2017; Guo et al. 2023), whereas no comprehensive transcriptome analysis has been conducted for the Japanese giant salamander. Consequently, current genetic assessments of the Japanese giant salamander still rely largely on small-scale sequence datasets.

Urodele amphibians generally possess extremely large genomes, often reaching several tens of gigabase (Gb). Among them, the giant salamanders are characterized by a primitive karyotype (Au type) with 2n = 60 chromosomes (Morescalchi et al. 1977; Morescalchi 1979). The estimated genome size of the giant salamander is approximately 50 Gb—about 16 times larger than that of humans—and has not yet been fully sequenced.

Within urodeles, the axolotl (Ambystoma mexicanum) is one of the few species for which a reference genome has been assembled, with a reported genome size of approximately 27 Gb (Smith et al. 2019). The axolotl possesses a different karyotype (S type) with 2n = 28 chromosomes.

In this study, we used the axolotl genome as a reference framework, as it represents the closest related species with an available assembled genome. By mapping giant salamander transcripts onto the axolotl genome, we constructed a putative chromosomal architecture for the giant salamander. Using this virtual chromosomal framework, polymorphisms identified in hybrids with the Chinese species were mapped to infer their approximate chromosomal positions and to estimate haplotype block structures. Elucidation of haplotype block distributions would enable reconstruction of hybridization history, provide insights into potential sex chromosome regions, and facilitate genome-wide association studies (GWAS) aimed at identifying genetic loci underlying morphological and behavioral traits.

In this study, we conducted transcriptome analyses of the Japanese giant salamander and hybrid individuals and performed virtual chromosomal mapping to comprehensively investigate the current genetic status of hybrid populations. This genome-wide framework provides a foundation for resolving the evolutionary dynamics and genetic consequences of hybridization in giant salamanders.

## Results

### RNA-seq Analysis of Japanese Giant Salamanders and Hybrids

To characterize the transcriptomes of Japanese giant salamanders and hybrids obtained from the Nabari region, we purified RNA from the skin of nine Japanese giant salamanders (three from Nichinan and six from Hanzaki-Lab) and twenty hybrids from Nabari, and performed RNA-seq analysis (Table 1, Fig. 1). Approximately 50 million paired-end reads per sample were generated (Table S1, Fig. 1). These reads were mapped to the Chinese giant salamander reference transcriptome, which consists of 30,366 transcripts (Geng et al. 2017). Because the sequence reads from all individuals were mapped to the reference transcriptome with high efficiency and without noticeable bias, it was demonstrated that polymorphism analyses of the Japanese giant salamander, the Chinese giant salamander, and their hybrids can be appropriately conducted using the Chinese giant salamander reference transcriptome (Table S2).

**Figure 1.**
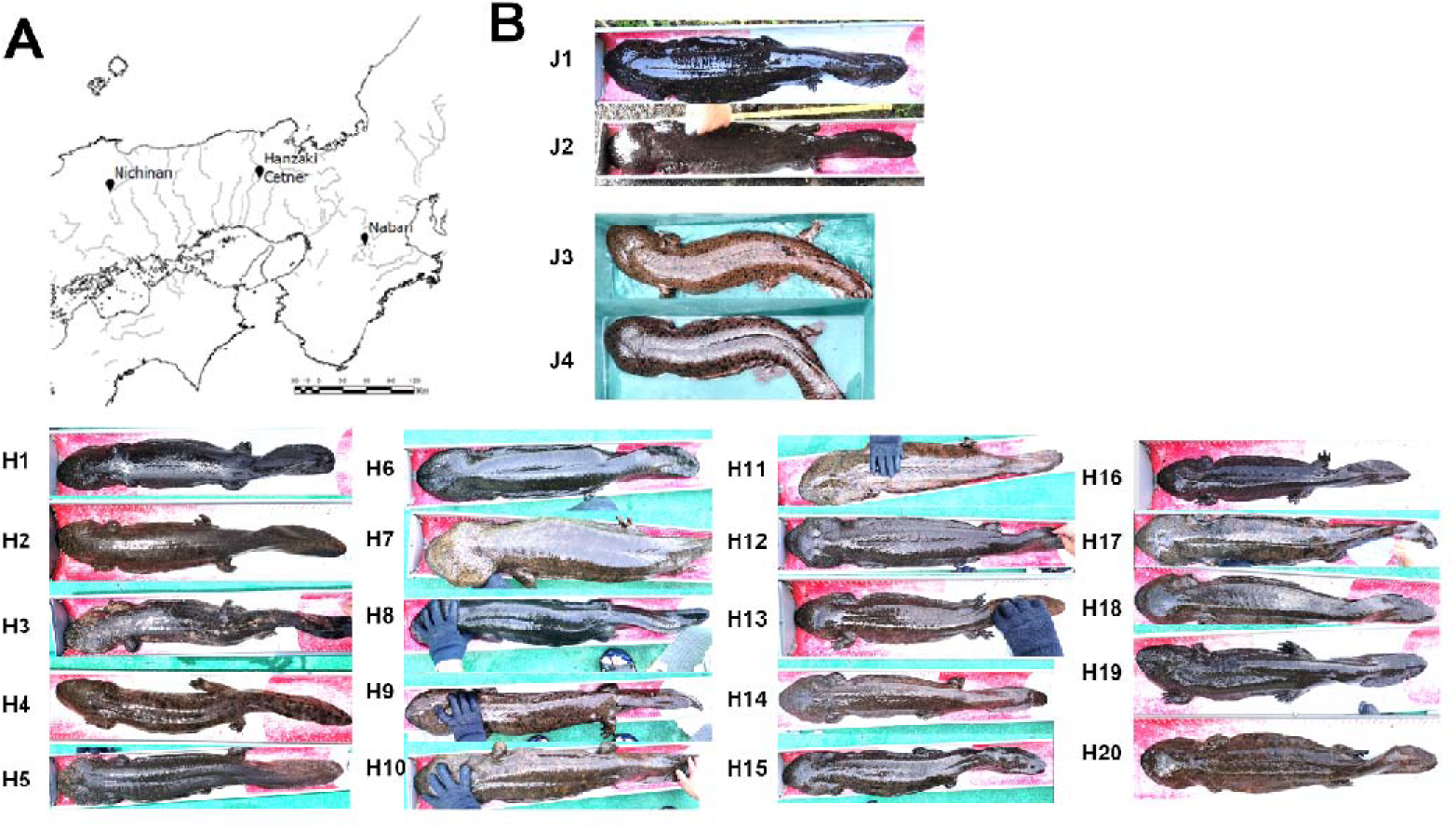
(A) Collection sites of Japanese giant salamander individuals, including Nichinan Town, Asago City (Hanzaki Laboratory), and Nabari City. (B) Dorsal views of Japanese giant salamanders and hybrid individuals identified in Nabari City. Japanese giant salamanders typically exhibit a dark brown to black coloration with fine mottling and numerous unpaired tubercles on the head. In contrast, Chinese giant salamanders are generally gray to beige in coloration, tend to have a flatter head and a shorter tail, and possess paired tubercles on the head.

**Table 1.**
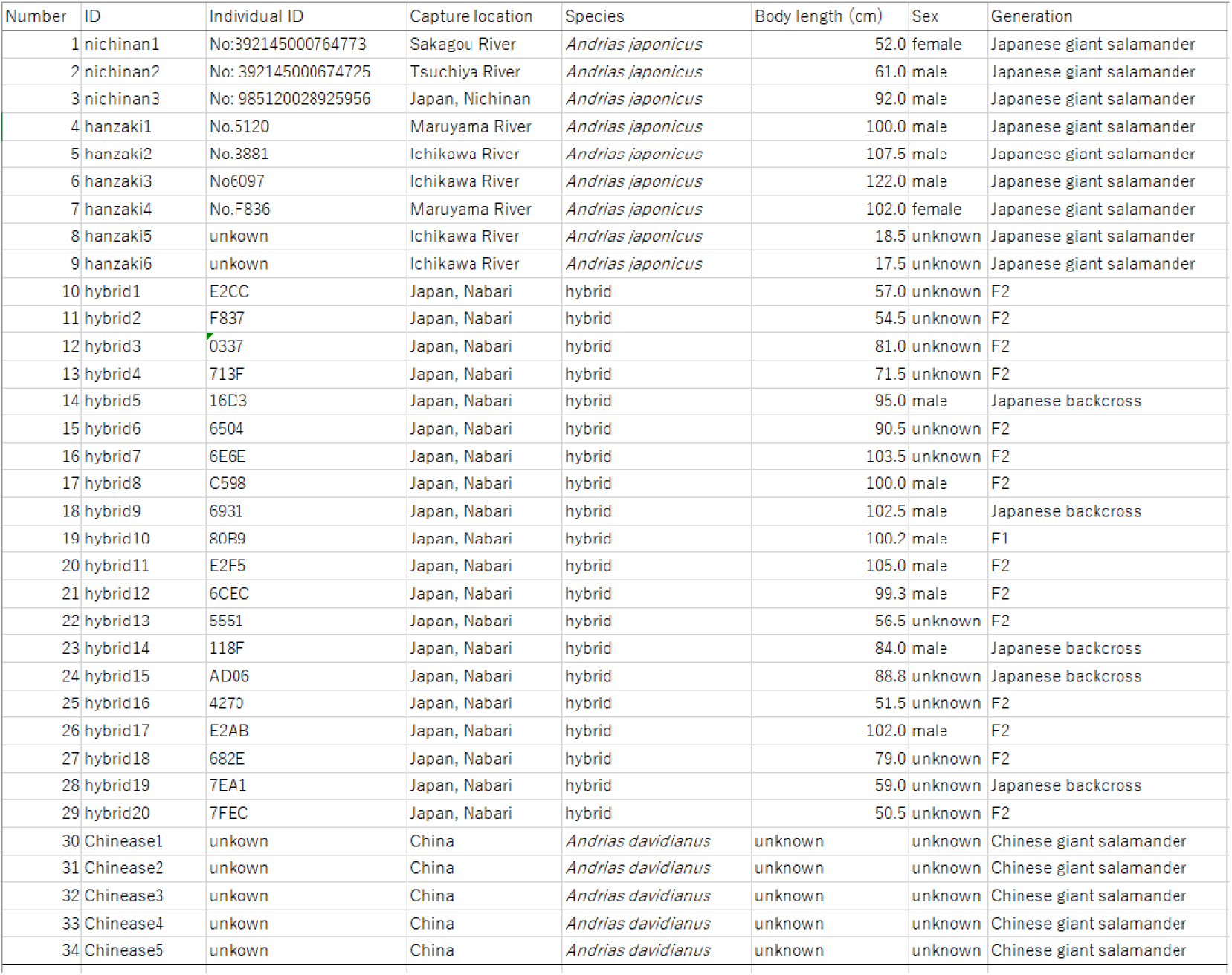
Individual data of the giant salamanders used in this study.

To enable comparison with Chinese giant salamanders, we also included previously reported RNA-seq data from five Chinese giant salamanders in this analysis (Geng et al. 2017; Bai et al. 2021). In total, we constructed a transcriptome dataset composed of nine Japanese giant salamander individuals, five Chinese giant salamander individuals, and twenty hybrid individuals (Table 1).

### SNP-based genotype analysis using the giant salamander transcriptome dataset

First, we performed sequence comparisons using transcriptome data from Japanese giant salamanders, hybrids, and Chinese giant salamanders. A total of 419,177 SNP candidates were identified as being specific to the Japanese giant salamander, the Chinese giant salamander, and the hybrid individuals, compared with the reference Chinese giant salamander genome (Table S3).

Next, we extracted SNPs located in highly expressed genes with sufficient sequence coverage. Gene expression levels were estimated using Kallisto (Table S4), and the top 5,000 genes with the highest TPM values were selected. To simplify subsequent analyses, we selected one representative SNP per gene based on the highest SNP quality score. To enable subsequent genomic synteny analysis using the axolotl genome sequence, we retained only genes annotated in the axolotl genome, resulting in a representative SNP dataset comprising 4,457 genes (Table S5). This dataset, including gene annotations, SNP information, and expression values, was used for the following analyses (Table S5). For example, when one gene (frmpd6) was visualized using the IGV browser, a putative SNP specific to the Japanese giant salamander was observed, and it was evident that some hybrid individuals possessed this SNP whereas others did not (Fig. S1). Thus, this dataset enables the investigation of gene-by-gene polymorphisms within hybrid individuals.

### PCA and NJ Tree Analyses of Transcriptome Datasets of Giant Salamanders

To assess the overall genetic variation across the 34 individuals, we performed principal component analysis (PCA) based on the transcriptome SNP dataset. The PCA identified four major clusters (Fig. 2A). As expected, Japanese giant salamanders and Chinese giant salamanders formed distinct and independent clusters.

**Figure 2.**
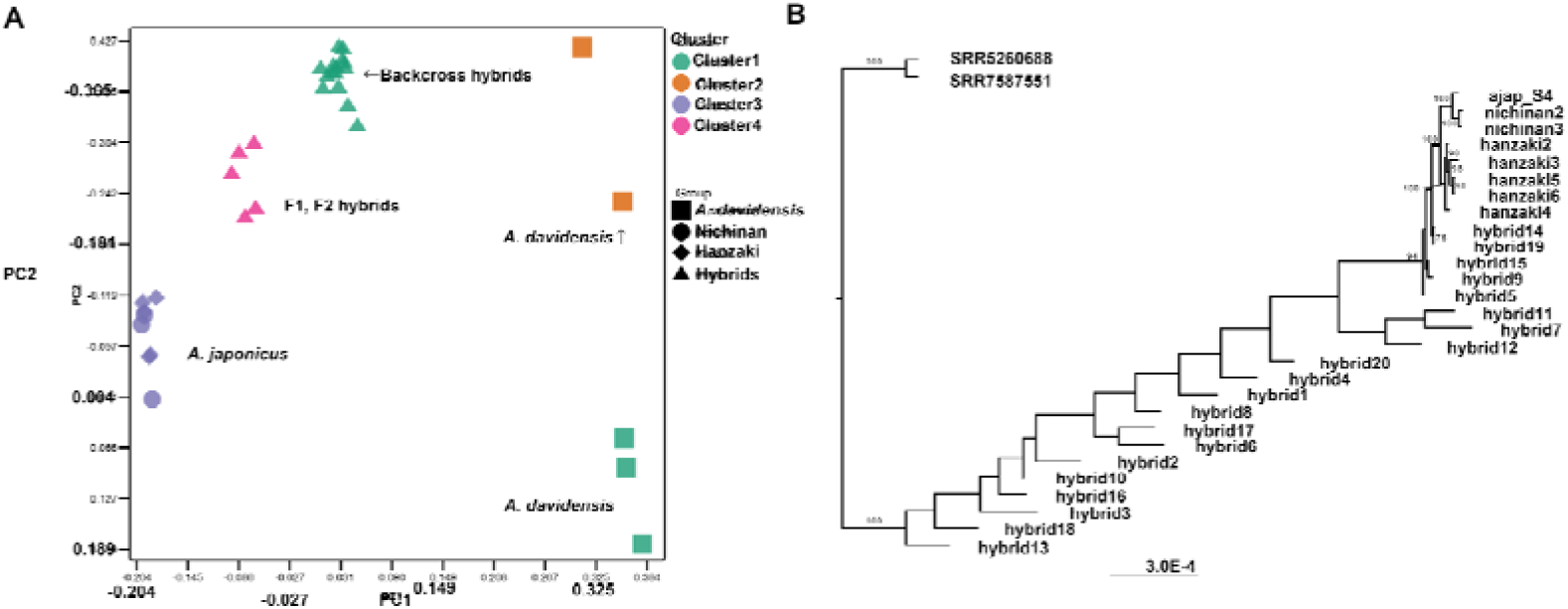
Gene expression analysis of giant salamander skin. (A) Principal component analysis (PCA) of transcriptome data from all 27 giant salamander individuals. The individuals were divided into four groups. (B) Neighbor-joining (NJ) phylogenetic tree of Japanese and Chinese giant salamanders constructed based on transcriptome data.

Interestingly, the 20 hybrid individuals were subdivided into two major groups. Five individuals (hybrid5, hybrid9, hybrid15, hybrid16, and hybrid19), designated as Group A, were positioned intermediate between the Japanese and Chinese giant salamander clusters. The remaining 15 individuals formed an independent cluster (Group B) and were located closer to the Chinese giant salamander population along the PC1 axis.

To perform a more detailed analysis, we conducted a neighbor-joining (NJ) tree analysis using 4,457 SNPs from the 34 individuals. In this analysis, the Chinese giant salamander population formed a distinct cluster and was clearly separated from the Japanese giant salamander cluster (Fig. 2B). The hybrids were divided into two groups: 15 individuals that clustered with the Chinese giant salamander lineage and five individuals that were more closely related to the Japanese giant salamander. The composition of these groups was consistent with the results of the PCA analysis. Taken together, both the PCA and NJ tree analyses revealed that the hybrid population was subdivided into two distinct groups.

### Genetic Characterization of the Two Hybrid Clusters

To identify the characteristics of the two hybrid groups, we performed further detailed analyses using the dataset of 34 individuals. All SNPs identified from the transcriptome analysis were genotyped and classified into three categories: putative Japanese homozygotes, Chinese homozygotes consistent with the reference sequence, and heterozygotes. The total number of SNPs in each category was calculated for each individual (Fig. 3). As expected, approximately 80% of the total SNPs in all five Chinese giant salamander individuals were classified as Chinese homozygotes. Similarly, approximately 90% of the total SNPs in all nine Japanese giant salamander individuals were classified as putative Japanese homozygotes. These results indicate that our dataset is sufficiently robust to distinguish the genetic backgrounds of the two species.

**Figure 3.**
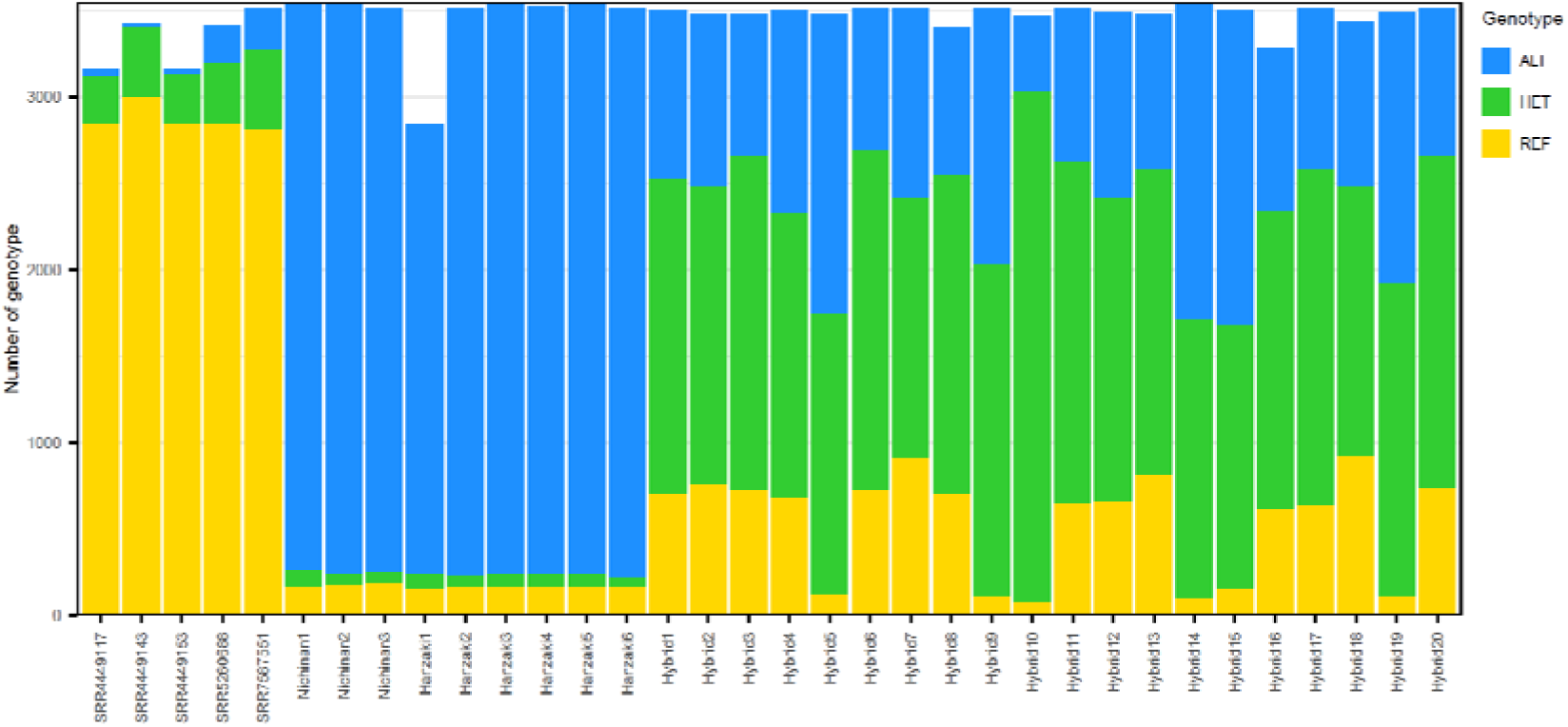
Overview of polymorphisms in the analyzed giant salamanders. SNPs were classified into three categories: Japanese homozygotes (yellow), Chinese homozygotes (blue), and heterozygotes (green). The number of SNPs in each category is shown for each individual. Hybrid individuals were further classified into three types: F1, F2, and backcross individuals (see Table S11).

We next investigated the hybridization history of the 20 hybrid individuals. Six individuals (hybrid5, hybrid9, hybrid10, hybrid14, hybrid15, and hybrid19) possessed relatively few Chinese homozygous SNPs (approximately 3% of total SNPs). Among these, only hybrid10 exhibited a high proportion of Japanese homozygous SNPs (81% of total SNPs), suggesting that this individual represents an F1 hybrid derived from a cross between Japanese and Chinese giant salamander parents. The remaining five individuals in this subgroup (hybrid5, hybrid9, hybrid14, hybrid15, and hybrid19) showed approximately 50% Japanese homozygous SNPs and 50% heterozygous SNPs, consistent with backcross individuals derived from crosses between F1 hybrids and Japanese giant salamanders.

The other fourteen individuals (including hybrid1 and hybrid2) exhibited an approximate SNP ratio of Japanese homozygotes: heterozygotes: Chinese homozygotes of 1:2:1. This segregation pattern suggests that these individuals most likely represent F2 progeny derived from intracrosses between F1 hybrids.

### Analysis of Transcripts Derived from the Mitochondrial Genome

Analysis of mitochondrial genetic variation in hybrid individuals provides insights into the maternal lineage and the sex-specific history of hybridization. Transcriptome data generated by RNA-seq include transcripts derived from the mitochondrial genome that are expressed in the sampled tissues, allowing analysis based on 13 protein-coding genes. We analyzed the mitochondrial genome origins of Japanese giant salamanders and hybrid individuals using alignment and variant-calling software. Previously reported mitochondrial reference sequences were used as genetic references (Zhang et al. 2003). Sequence analyses were conducted for the 13 protein-coding genes encoded in the mitochondrial genome, and polymorphisms were identified for all individuals (Fig. 4). Interestingly, all hybrid individuals possessed mitochondrial haplotypes of Japanese origin. This result indicates that, if hybridization occurred within two generations, the mother or maternal grandmother of the hybrid individuals was of Japanese origin.

**Figure 4.**
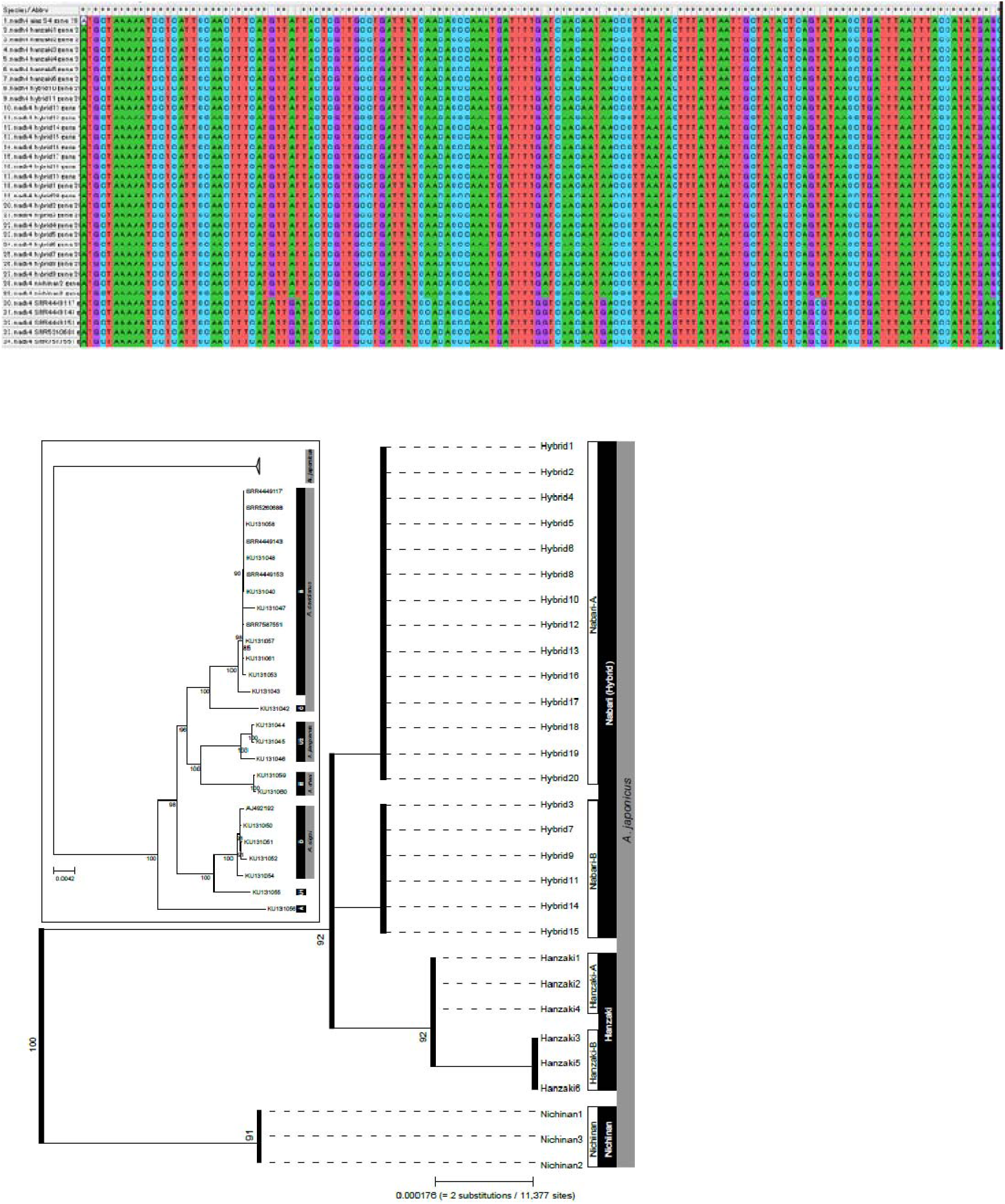
Analysis of mitochondrial genome-encoded genes in hybrid giant salamanders. (Upper panel) Nucleotide alignment of partial *ND4* sequences and their polymorphic sites, visualized using MEGA12 software. **(Middle panel)** A maximum-likelihood phylogenetic tree reconstructed from the nucleotide sequences of 13 mitochondrial protein-coding genes using IQ-TREE v3.0.1. **(Lower panel)** The clade containing Japanese giant salamanders and hybrid individuals is shown in a compressed form. Bootstrap support values (%) are indicated at nodes with values greater than 90.

### Analysis of Differentially Expressed Genes in the Skin of the Japanese Giant Salamander and Its Hybrids

To evaluate the effects of hybridization on skin tissue, we analyzed genes exhibiting differential expression between the Japanese giant salamander and hybrid individuals. Read counts mapped to 30,366 reference genes were quantified gene expression levels. Differentially expressed genes (DEGs) were identified by comparing nine Japanese giant salamander individuals with twenty hybrid individuals. A total of 1,343 genes were identified as significantly upregulated in hybrids compared with the Japanese giant salamander (Table S6). Gene Ontology (GO) enrichment analysis of these genes revealed significant enrichment in 214 GO terms, including response to toxic substance (GO:0009636), response to temperature stimulus (GO:0009266), and adaptive immune response (GO:0002250) (Table S7). In contrast, 1,050 genes were found to be significantly downregulated in hybrids relative to the Japanese giant salamander (Table S6). GO enrichment analysis of these genes identified significant enrichment in 97 GO terms, including regulation of innate immune response (GO:0045088) and response to virus (GO:0009615) (Table S9).

We further examined which specific genes exhibited differential expression in each individual. In the skin of hybrids, increased expression was observed for genes involved in oxidative stress responses, including SODC5 (superoxide dismutase [Cu-Zn] 5), CATA (catalase), and THIO (thioredoxin); heat shock proteins including HSP70/90 family members (HSP7C, heat shock protein 70 family; HS90A, heat shock protein HSP90-alpha) and the HSP40 family (DNJC3, DnaJ homolog subfamily C member 3; DNJA4); regulation of reactive oxygen species (ROS) production (DUOX2, dual oxidase 2); xenobiotic metabolism (ADH4, alcohol dehydrogenase 4); and genes related to neural responses to stimuli such as temperature (ADRB2, adrenoceptor beta 2; CHRNA7 (ACHA7), cholinergic receptor nicotinic alpha 7 subunit) (Table S8).

In contrast, in the Japanese giant salamander group, increased expression was observed for genes associated with activation of antiviral responses, including CCL4 (C-C motif chemokine ligand 4), CCL5, CCL19, CXL10 (C-X-C motif chemokine ligand 10), OAS3 (2’-5’-oligoadenylate synthetase 3), OASL2 (2’-5’-oligoadenylate synthetase-like 2), and IRF1 (interferon regulatory factor 1) (Table S10). A detailed discussion of these genes is provided below.

### Construction of a Putative Genome Structure of the Japanese Giant Salamander Based on Axolotl Genome Synteny Analysis

Although analysis of total SNP genotypes in hybrids provided information regarding their hybridization history, we anticipated that genome-wide analysis of haplotype blocks would enable a more detailed reconstruction of this history. If a whole-genome assembly of the giant salamander were available, the 4,457 genes identified in this study could serve as markers to infer haplotypes genome-wide. However, to date, no whole-genome assembly of the giant salamander has been reported. Therefore, we constructed a putative synteny map of the giant salamander genome based on the genome sequence of the axolotl (*Ambystoma mexicanum*), a urodele species whose whole genome has been sequenced (based on NCBI genome data; Table S13). Using this synteny information, we generated a pseudo genome-wide polymorphism map of the giant salamander (Fig. 5; Table S11).

**Figure 5.**
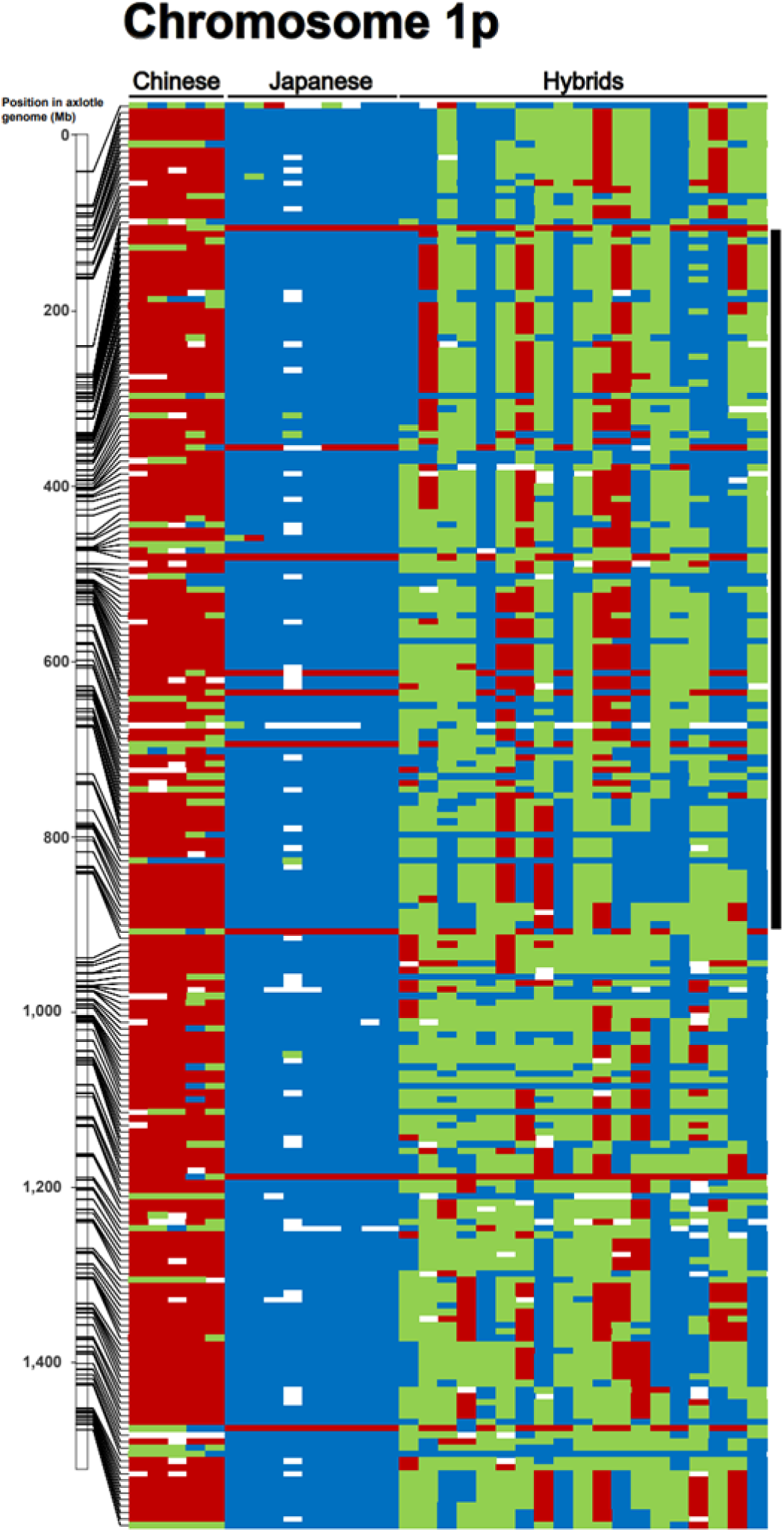
Genome-wide distribution of SNPs from 27 giant salamander individuals mapped onto the axolotl genome based on syntenic relationships (chromosome 1p shown as an example). Haplotype blocks are indicated by black bars on the left. Chinese giant salamander, Japanese giant salamander, and hybrid individuals are shown on the x-axis, whereas base-pair positions along the syntenic chromosome are shown on the y-axis. Chinese homozygotes (red), Japanese homozygotes (blue), and heterozygotes (green) are indicated by different colors

Because these genes were preselected based on their known synteny with the axolotl, all 4,457 genes were successfully mapped onto chromosomes, and virtual chromosome numbers and gene positions were assigned (Table S11, Fig. S2). On average, approximately 300 genes were mapped per chromosome across the 14 virtual chromosomes corresponding to axolotl genome. As expected, long haplotype blocks ranging from approximately 100 Mbp to 600 Mbp were observed in the hybrid individuals. In many individuals, several haplotype blocks were detected per chromosome. This pattern is consistent with the hybrids being F1, F2, or backcross individuals. For example, in hybrid9, a Japanese homozygous haplotype block spanning approximately 570 Mbp was observed on chromosome 1p (NC_090921.1), from position 274,182,923 to 839,508,182 (Fig. 5). In the same chromosomal region, hybrid2 exhibited a Chinese homozygous haplotype block spanning approximately 160 Mbp, from position 271,344,831 to 432,983,236 on chromosome 1p (Fig. 5).

In this manner, haplotype blocks were observed across all chromosomes. The regions in which these haplotype blocks are maintained indicate conserved genomic segments between the axolotl and the giant salamander. At the same time, the presence of such long haplotype blocks suggests that the number of hybrid generations is still limited and that extensive recombination has not yet occurred. In contrast, some regions lacked clearly defined long haplotype blocks. For example, on chromosome 1p, from position 841,682,162 to 1,202,677,982 (approximately 360 Mbp), relatively short haplotype blocks of several tens of megabases were interspersed. This region may represent a portion of the genome where synteny between the axolotl and the giant salamander is less well conserved. In addition, several polymorphisms that were specific to the Chinese giant salamander individuals, the Japanese giant salamander individuals, or the hybrid individuals analyzed in this study were observed. However, such regions constitute only a limited fraction of the genome overall. In many other regions, the virtual chromosome map appears to be highly informative and useful.

Based on the virtual chromosome information of the giant salamander, we investigated whether environmental adaptation–related genes—such as immune response genes including MHC (Major Histocompatibility Complex) (Kiemnec-Tyburczy et al. 2012), Toll-like receptors, TLR3, TLR4, and TLR5 (Roach et al. 2005); temperature adaptation genes TRPA1 and TRPV1 (Saito and Tominaga 2017; Saito et al. 2022); and the pigmentation-related gene MC1R (Melanocortin 1 receptor) (Li et al. 2025)—exhibited biased patterns of genetic polymorphism in the hybrid population. We examined the haplotypes at these seven loci. For all loci and their surrounding regions, multiple individuals among the 20 hybrids were identified as Japanese homozygotes, Chinese homozygotes, or heterozygotes, and no obvious genotype bias was detected (Table S12). At least for these genes, it was therefore evident that no clear skew in genotype frequencies was present in this hybrid population, which consists of individuals with a relatively small number of hybrid generations.

## Discussion

### Hybridization history and the origin of introduced species

To date, comparisons between the Japanese giant salamander and the Chinese giant salamander have primarily focused on mitochondrial genomes and a limited number of genetic markers (Yoshikawa et al. 2011; Yoshikawa et al. 2012; Turvey et al. 2019). In this study, we report for the first time genome-wide SNP variation in the Japanese giant salamander based on transcriptome analysis. Furthermore, we analyzed SNPs from three geographically distinct populations of the Japanese giant salamander and identified region-specific SNPs in each population. These findings are expected to serve as a fundamental resource for future studies on the evolution, divergence, and conservation of the Japanese giant salamander.

Furthermore, SNP analysis of 20 hybrid individuals revealed that F1 hybrids were extremely rare, whereas F2 individuals were the most abundant, followed by backcrosses. In addition, analysis of predicted haplotype blocks based on constructed pseudochromosomes suggested that the likelihood of hybridization beyond the F2 generation is low. The absence of backcrossing with the Chinese giant salamander suggests that the initial number of introduced Chinese giant salamander individuals was likely small.

Furthermore, mitochondrial genome analysis suggests that these initial Chinese giant salamander individuals were likely males. Although estimating the generation time of giant salamanders in the wild is challenging, growth data indicate that it is approximately 15–20 years (Yamasaki et al. 2017). The hybrid individuals analyzed in this study were large in size and are estimated to be over 30 years old; therefore, the introduction of the alien species likely occurred more than 50 years ago (Fig. 6). This timing is consistent with the period when commercial importation from China to Japan began in the 1970s (Takakura et al. 2025).

**Figure 6.**
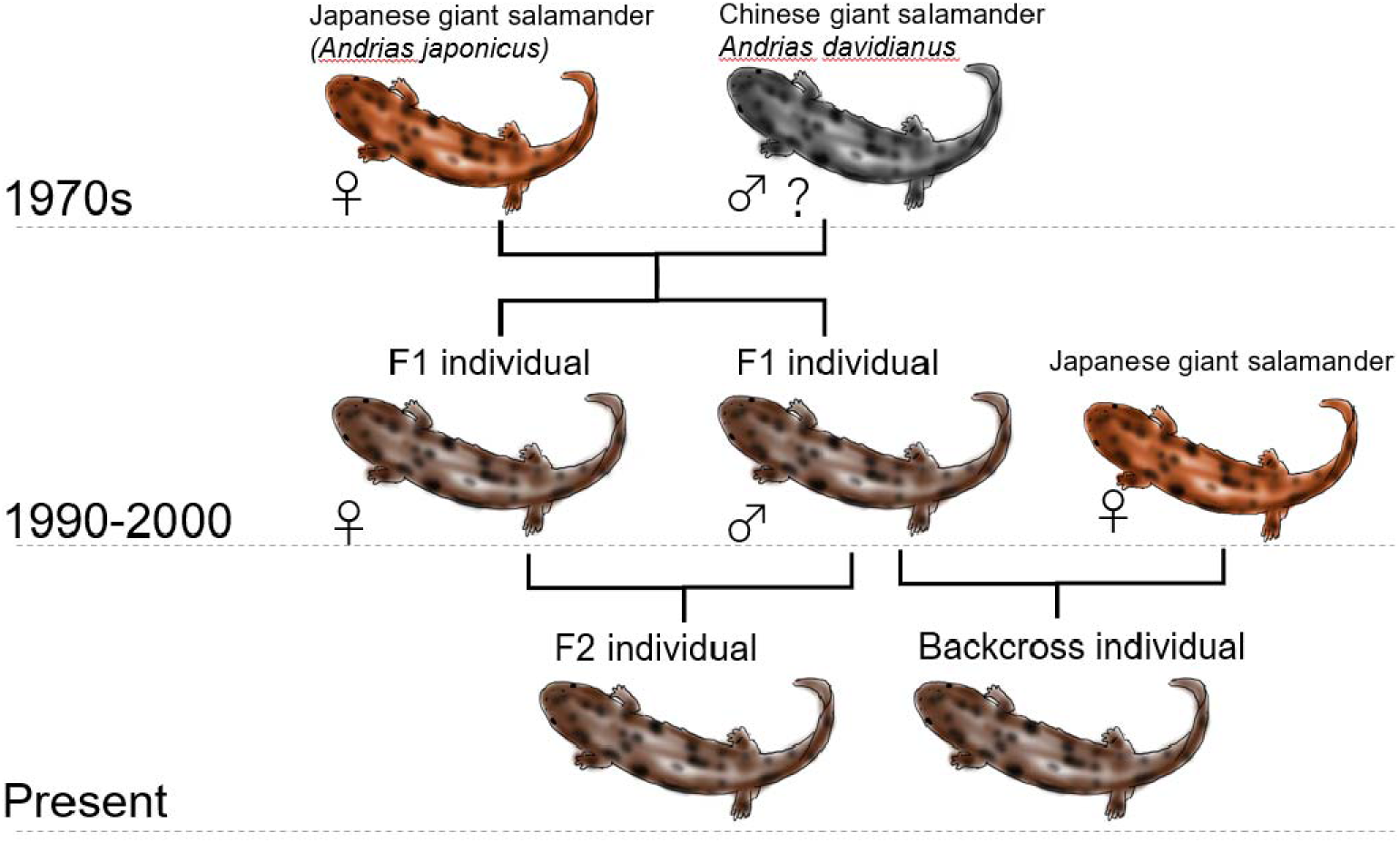
Inferred history of hybridization among giant salamander populations. Mitochondrial genome analysis suggests that the initially introduced Chinese giant salamanders were likely males. Although it is difficult to estimate the generation time of giant salamanders in the wild, available growth data suggest a generation time of approximately 15–20 years. The hybrid individuals analyzed in this study were large-bodied and are estimated to be more than 30 years old. Therefore, the introduction of Chinese giant salamanders into the study area likely occurred more than 50 years ago. This estimate is consistent with the period when commercial importation of Chinese giant salamanders into Japan began in the 1970s.

Sexually mature, large male giant salamanders construct nesting burrows along riverbanks. Females enter these burrows to mate and lay eggs (Okada et al. 2015); however, males provide parental care for the eggs and larvae (Takahashi et al. 2017). Because multiple females may visit and spawn within a single nest, genes derived from a particular male can spread efficiently within a population. In the present study, all hybrid individuals were found to possess mitochondrial DNA of Japanese origin. This finding indicates that male Chinese giant salamanders have contributed to reproduction within these hybrid populations. If male reproductive success is higher in Chinese giant salamanders or hybrids than in pure Japanese giant salamanders, the spread of hybrid lineages could proceed rapidly. Such a scenario raises serious concerns for environmental conservation and the persistence of the native species. Therefore, urgent management actions, including the isolation and control of hybrid individuals, are strongly warranted.

### Functional characterization of differentially expressed genes and environmental responses

In the skin, we identified 1,343 genes that were upregulated in hybrids and 1,050 genes that were downregulated compared with the Japanese giant salamander. Functional analysis of these differentially expressed genes revealed that the genes upregulated in hybrids are involved in oxidative stress responses (SODC5, CATA, THIO) (Zelko et al. 2002; Lu and Holmgren 2014; Glorieux and Calderon 2017), regulation of reactive oxygen species (ROS) production (DUOX2) (Bedard and Krause 2007), and xenobiotic metabolism (ADH4) (Vasiliou et al. 2000). These findings suggest the activation of an integrated oxidative stress response network, which is strongly associated with inflammation control and responses to ultraviolet (UV) radiation (Sies 2015). The simultaneous induction of HSP70/90 and HSP40 family genes observed in hybrids likely reflects the activation of protein homeostasis (proteostasis) mechanisms (Kampinga and Craig 2010; Taipale et al. 2010). Furthermore, changes in ADRB2 and CHRNA7 support the possibility that neural stimuli, such as temperature, regulate tissue responses via immunomodulatory pathways (Wang et al. 2003; Sanders 2012). The observed upregulation of these gene sets in hybrids suggests that excessive stress responses are induced in the skin of hybrid individuals. One possible explanation is that hybrids are less well adapted to the environmental conditions of Japanese rivers than pure native individuals, and therefore cope with these conditions by activating stress response pathways.

In contrast, genes upregulated in the Japanese giant salamander, including CCL5, CXCL10, CCL4, CCL19, OASL2, OAS3, and IRF1, suggest activation of a type I interferon-dependent antiviral response (Schneider et al. 2014). Furthermore, the induction of chemokine genes indicates activation of defense mechanisms through the recruitment of immune cells to skin tissues (Appay and Rowland-Jones 2001). The observed downregulation of these gene sets in hybrids suggests that immune responses to microorganisms may differ between hybrid individuals and pure native individuals.

In summary, these gene sets have the potential to serve as markers for distinguishing hybrid individuals based on their expression levels. However, because the Japanese giant salamander and hybrid individuals analyzed in this study were collected from different locations, it is also possible that the observed gene expression differences reflect variation in their environmental conditions. Nevertheless, our results demonstrate that these gene sets exhibit clear expression changes in the skin, suggesting that they may represent a useful marker set for reflecting hybridization status and habitat conditions in giant salamanders. Future studies incorporating a wider range of hybrid and pure populations, as well as samples from diverse locations, are expected to enable more accurate analyses.

### The hypothetical genome map of giant salamander

Giant salamanders produce approximately 300–700 eggs per breeding event; however, in the wild, only a very small number of individuals survive to adulthood. Among hybrid individuals that successfully reach maturity, it is expected that genotypes may be biased toward either the Japanese or Chinese lineage in genes conferring survival advantages, such as those involved in immune responses (e.g., MHC and TLR) and environmental sensitivity factors (e.g., temperature tolerance and toxin metabolism). On the other hand, to investigate whether hybridization leads to biased genotypes in specific genes, it is desirable to examine the genotypes of as many genes as possible. However, in the present study, genotypes were determined only for 4,457 genes that are highly expressed in the skin. Considering that the total number of genes in amphibians is estimated to be on the order of twenty thousand, this coverage is not sufficient. Nevertheless, genes located in close proximity on chromosomes are likely to share the same genotype due to the hitchhiking effect, forming haplotype blocks. If the whole-genome sequence of the giant salamander were available, it would be possible to use the genotypes of the 4,457 genes obtained in this study as markers to infer the haplotypes of nearly all genes. However, to date, the whole-genome sequence of the giant salamander has not been determined. Therefore, we constructed a synteny map of the giant salamander based on the genome sequence of the axolotl, a urodele species with a fully sequenced genome, and generated a hypothetical genome-wide polymorphism map of the giant salamander. We then performed synteny mapping of the 4,457 giant salamander genes onto the axolotl genome. The Japanese giant salamander has a primitive chromosome type (Au) with 2n = 60. In contrast, the axolotl has a type (S) with 2n = 28 (Morescalchi 1979). Since changes in chromosome number are mainly caused by chromosomal fusion, it is likely that majority of the gene synteny within chromosomes is conserved (Smith et al. 2019). In this study, although the synteny map between giant salamander and axolotl likely partially contains regions of low reliability, particularly in areas where chromosomal structural changes such as chromosome junctions have occurred. However, the overall arrangement of gene synteny is considered to be broadly reliable, since based on the size of haplotype blocks, synteny spanning several tens to hundreds of megabase (Mb) appears to be conserved. This hypothetical genome map is expected to become a powerful tool for identifying associations between genes and single nucleotide variants (SNVs) underlying giant salamander phenotypes, including morphological and ecological traits, through GWAS analyses. Furthermore, these insights are anticipated to serve as an important resource for elucidating the evolution of the giant salamander and the molecular mechanisms underlying species-specific traits.

In this study, the analysis was limited due to the restricted dataset of only 20 individuals. However, by increasing the sample size to hundreds of individuals in future studies, it may become possible to identify genetic polymorphisms associated with environmental adaptation through hybridization, as well as those controlling morphological and coloration differences between the Chinese and Japanese giant salamander species. Furthermore, by identifying gene sets responsible for the characteristic body shape and coloration of non-native species, it is expected that criteria for distinguishing invasive individuals can be established, thereby contributing to species conservation and environmental management. The genome information obtained in this study for the Japanese and Chinese giant salamanders is therefore expected to serve as a fundamental resource for future research and conservation activities on giant salamanders.

## Material and Methods

### Animals

A total of nine Japanese giant salamanders (three from Nichinan and six from Hanzaki Lab) and twenty hybrids from Nabari were used in this study. Skin tissue samples were collected under the following permits: Permit No. 202200103090 (Tottori Prefectural Directive) and Asago City Document No. 289 (Choubun), dated January 5, 2023, and Nabari City Board of Education Document No. 513, dated July 17, 2024.

### Purification of RNA and RNA-seq analysis

Individuals of the giant salamander were anesthetized under low-temperature conditions, and approximately 0.5 g of tissue was collected from the tail fin. Total RNA was extracted using TRIzol (Invitrogen) according to the acid guanidinium thiocyanate–phenol–chloroform (AGPC) method following the manufacturer’s instructions. RNA concentration was measured using a NanoDrop spectrophotometer (Invitrogen). RNA quality was assessed by agarose gel electrophoresis and ethidium bromide staining, and the presence of 18S and 16S ribosomal RNA bands was confirmed. The RNA samples were subsequently analyzed by Macrogen Japan, Inc., Tokyo, Japan. Transcriptome resequencing was performed using paired-end reads (151 bp) on a NovaSeq X platform. Libraries were prepared using the TruSeq Stranded mRNA Library Preparation Kit according to the manufacturer’s protocol.

### Mitochondrial gene assembling and nuclear gene genotyping using RNA-seq Data

To reconstruct nucleotide sequences from each sample, RNA-seq reads were used for assembling of mitochondrial genes and genotyping of nuclear genes. In addition to the RNA-seq data generated in this study, publicly available RNA-seq data from pure Chinese giant salamanders (Geng et al., 2017; Bai et al., 2021) were retrieved from the SRA database (SRR4449117, SRR4449143, SRR4449153, SRR5260688, and SRR7587551). Raw sequencing reads were quality-checked using FastQC v0.12.0 and trimmed with Trimmomatic v0.39 to remove adapter sequences and low-quality bases.

Mitochondrial genome sequences were assembled from quality-filtered RNA-seq reads using the MITOGARD pipeline (Nachtigall et al. 2021). MITOGARD recruits mitochondrial reads by mapping them against a phylogenetically related reference mitochondrial sequence, and then assembles the recruited reads to reconstruct mitochondrial genome sequences. In this study, a known mitochondrial genome sequence of Chinese giant salamander (Zhang et al., 2003) was used as the reference. The recruited reads were assembled into contigs using the Trinity assembler. The resulting mitochondrial contigs were automatically annotated using the MITOS web server (10.1016/j.ympev.2012.08.023).

For nuclear genes, the filtered reads were aligned to the reference transcriptome of Chine giant salamander (Geng et al. 2017) using STAR v2.7.11b (Dobin et al., 2013) with the parameter --outFilterMultimapNmax 1. Genotyping was performed using Genome Analysis Toolkit (GATK) v4.6.1.0 (McKenna et al., 2010; Van der Auwera et al., 2013) following GATK Best Practices for RNA-seq variant calling. Read groups were added using samtools addreplacerg, and duplicate reads were marked using GATK MarkDuplicates. The SplitNCigarReads tool was applied to split reads spanning splice junctions and to hard-clip overhanging sequences in intronic regions.

Variants were called for each individual separately using HaplotypeCaller in RNA-seq mode (--emitRefConfidence GVCF), generating GVCF files. Individual GVCF files were combined using CombineGVCFs, followed by joint genotyping with GenotypeGVCFs to produce a multi-sample VCF file. Variants were filtered using VariantFiltration with the following thresholds: QD < 2.0, FS > 200.0, and SOR > 10.0.

### Phylogenetic analyses based on mitochondrial and nuclear genes

We reconstructed phylogenetic relationships of the samples used in this study based on mitochondrial and nuclear genes. For analyses of the 13 protein-encoding mitochondrial genes, multiple nucleotide sequence alignment was performed using MUSCLE (Edgar 2004) as implemented in MEGA version 12 (Tamura et al. 2021). Maximum likelihood (ML) phylogenetic trees were inferred using IQ-TREE version 3.0.1 (Álvarez-Carretero et al. 2023), with the best-fit substitution model selected by the built-in ModelFinder (Gossmann et al. 2016) based on the Bayesian Information Criterion (BIC). Branch support was assessed with 1,000 ultrafast bootstrap replicates (Hoang et al. 2018).

For nuclear gene phylogenetic analysis, we first identified genes where all regions were covered by at least 10 reads for each sample, which yielded approximately 2,000–3,000 qualifying genes for most samples. However, four samples (hanzaki1, SRR4449117A, SRR4449143A, and SRR4449153A) exhibited substantially lower coverage with only approximately 1,000 qualifying genes each and were therefore excluded from subsequent analyses to ensure data quality and consistency. From the remaining samples, we extracted genes where all regions were covered by at least 10 reads across all samples, resulting in a final dataset of 1,350 loci. FASTA sequences for these loci were generated from VCF files using bcftools consensus (10.1093/bioinformatics/btp352), with IUPAC ambiguity codes applied to represent heterozygous sites and without phasing. Phylogenetic inference was performed using IQ-TREE version 3.0.1 (Álvarez-Carretero et al. 2023) with the best-fit substitution model selected by ModelFinder (Gossmann et al. 2016) based on BIC, and maximum likelihood trees were constructed with branch support assessed using 1,000 ultrafast bootstrap replicates (Hoang et al. 2018).

### Transcript abundance quantification and differential gene expression (DEG) analysis

Transcript abundance was quantified using Kallisto v0.46.2 (Bray et al., 2016; doi:10.1038/nbt.3519) with pseudo-alignment against the reference transcriptome. A Kallisto index was built using the default k-mer size (k = 31), and paired-end reads were quantified with 100 bootstrap replicates to estimate technical variance. Transcript-level TPM values were summarized to gene-level expression using tximport (Soneson et al. 2015). Genes with TPM ≥ 1 in at least one sample were retained for downstream analyses to focus on genes expressed in skin tissue.

Differential gene expression (DEG) analysis was performed using Sleuth (version 0.30.1) (Pimentel et al. 2017), an R package designed to work with Kallisto output. Sample metadata, including genetic background (pure or hybrid) and locality, were imported from a metadata file. A Sleuth object was constructed using the sleuth_prep() function with the experimental design formula ∼condition, which normalizes transcript-level estimates and aggregates them to the gene level. The full model incorporated the condition variable to test for differential expression between experimental groups.

To identify differentially expressed genes, a likelihood ratio test (LRT) approach was employed. Two models were fitted to the data: a full model (∼condition), which included the experimental condition as a predictor, and a reduced (intercept-only) model that excluded this variable. The sleuth_fit() function was used to fit both models, and sleuth_lrt() was applied to perform the likelihood ratio test comparing the reduced model with the full model. Differential expression results were extracted using sleuth_results() with the test type set to “lrt”. Genes were considered significantly differentially expressed if they met a false discovery rate (FDR) threshold of q-value < 0.05, as determined using the Benjamini–Hochberg correction for multiple testing.

### GO term enrichment analysis

Differentially expressed genes (DEGs) were uploaded to the Gene Ontology Consortium website (https://geneontology.org/) and analyzed using the Biological Process ontology. Because of its extensive functional annotation and the high reliability of experimentally validated biological information, *Mus musculus* was selected as the reference species for the enrichment analysis.

Among the genes upregulated in hybrids, 853 gene IDs were successfully mapped, and 214 GO terms were significantly enriched at an FDR-adjusted P value of < 0.05 (Table S7). Among the genes downregulated in hybrids, 777 gene IDs were successfully mapped, and 97 GO terms were significantly enriched at an FDR-adjusted P value of < 0.05 (Table S9).

### Cross-Species Synteny Mapping between the Giant Salamander Transcriptome and the Axolotl Genome

To identify syntenic relationships between the giant salamander and the axolotl, whose genome sequence is available, we performed sequence similarity searches using the Basic Local Alignment Search Tool (BLAST). We used the complete set of transcript sequences (transcriptome) of the Chinese giant salamander (Geng et al. 2017) and the predicted protein sequences (proteome) from the axolotl RefSeq genome assembly (Smith et al. 2019). Specifically, BLASTX searches were conducted using the Chinese giant salamander transcriptome as queries against the axolotl proteome database. Conversely, TBLASTN searches were performed using the axolotl proteome as queries against the Chinese giant salamander transcriptome database. Orthologous genes were identified based on reciprocal best hits (RBHs), which were defined as the best-matching sequences in both search directions and are presumed to represent genes that diverged through speciation. Of the 22,060 Chinese giant salamander transcripts analyzed, 13,708 were identified as reciprocal best hits.

## Data availability

The RNA-seq reads generated in this study are available in the NCBI database under BioProject accession number PRJNA1273478. The previously reported BioProject is available under accession number PRJNA433898.

## Version of software programs

An overview of the analysis pipeline and the versions of the software used are provided in Figure S3.

## Acknowledgments

The authors thank Karin Obata, Wakana Fujii, Chinatsu Tada, Akimasa Maki, Ryouga Okada,Chiyo Nambafor experimental support. We gratefully acknowledge Dr. Ikuo Miura (Hiroshima University), Dr. Kanto Nishikawa (Kyoto University), and Dr. Hajime Ogino (Hiroshima University) for their valuable discussions.

## Author contributions

T.I., and Y.O. designed the study. T.I., and Y.O. supervised the project. T.I., S.O., M.S., R.T., M.Y., Z.S., and Y.O. conducted the laboratory experiments. T.I., H.B., and Y.O. performed the bioinformatic analysis. T.I., H.B., and Y.O. wrote the manuscript. All authors provided feedback on manuscript.

