## Supplementary material for "Transcriptome-based genome-wide analysis reveals hybridization dynamics and genetic structure of Japanese giant salamanders": Supple Fig

**Supplemental Figures**

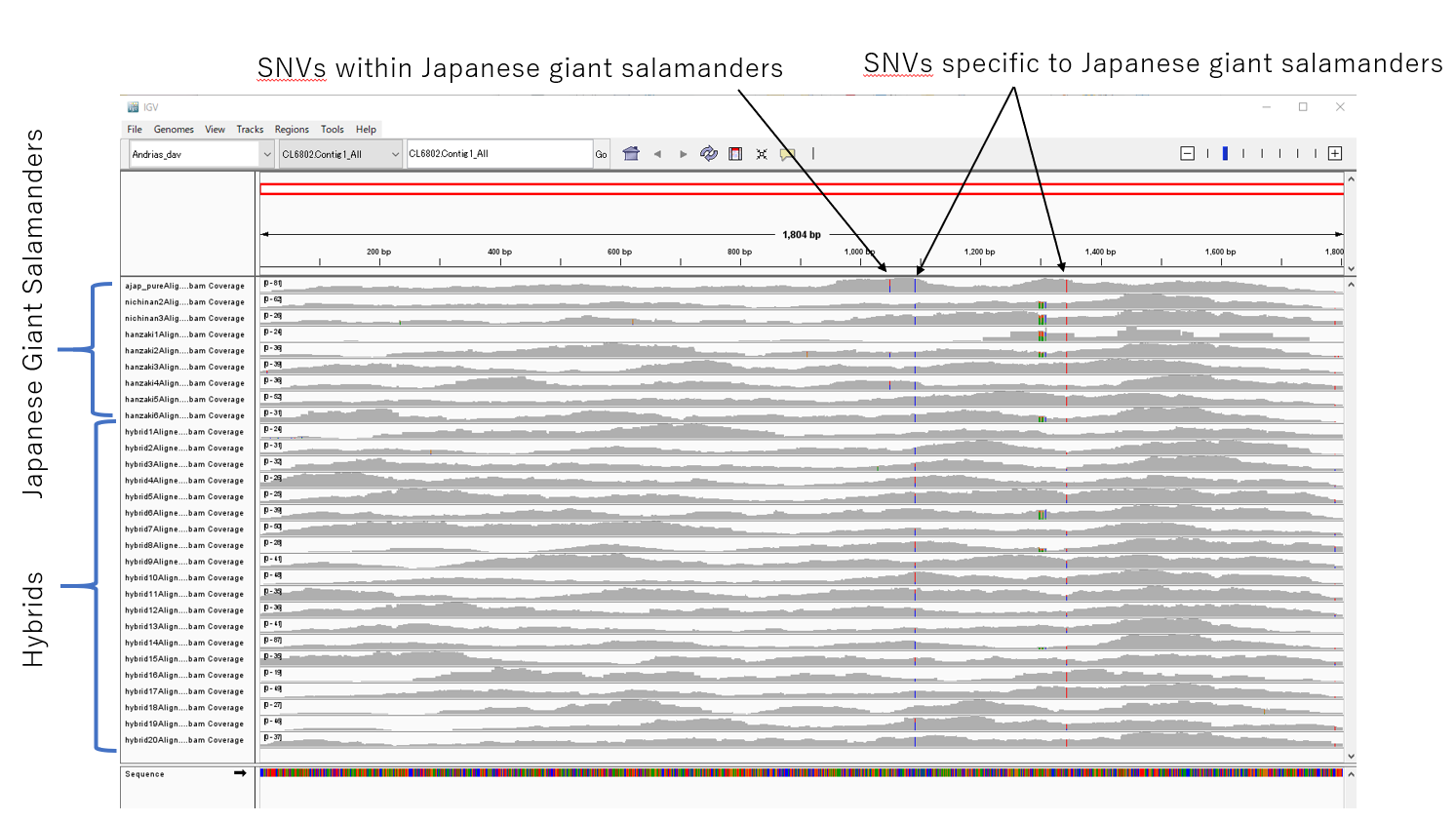

**Figure S1. Representative IGV view of the frmpd6 gene (ID: CL6802).** Mapped sequence data from nine Japanese giant salamanders and 20 hybrid individuals are shown in the left panel. One SNV polymorphic within Japanese giant salamanders and two SNVs specific to Japanese giant salamanders were identified (arrows).

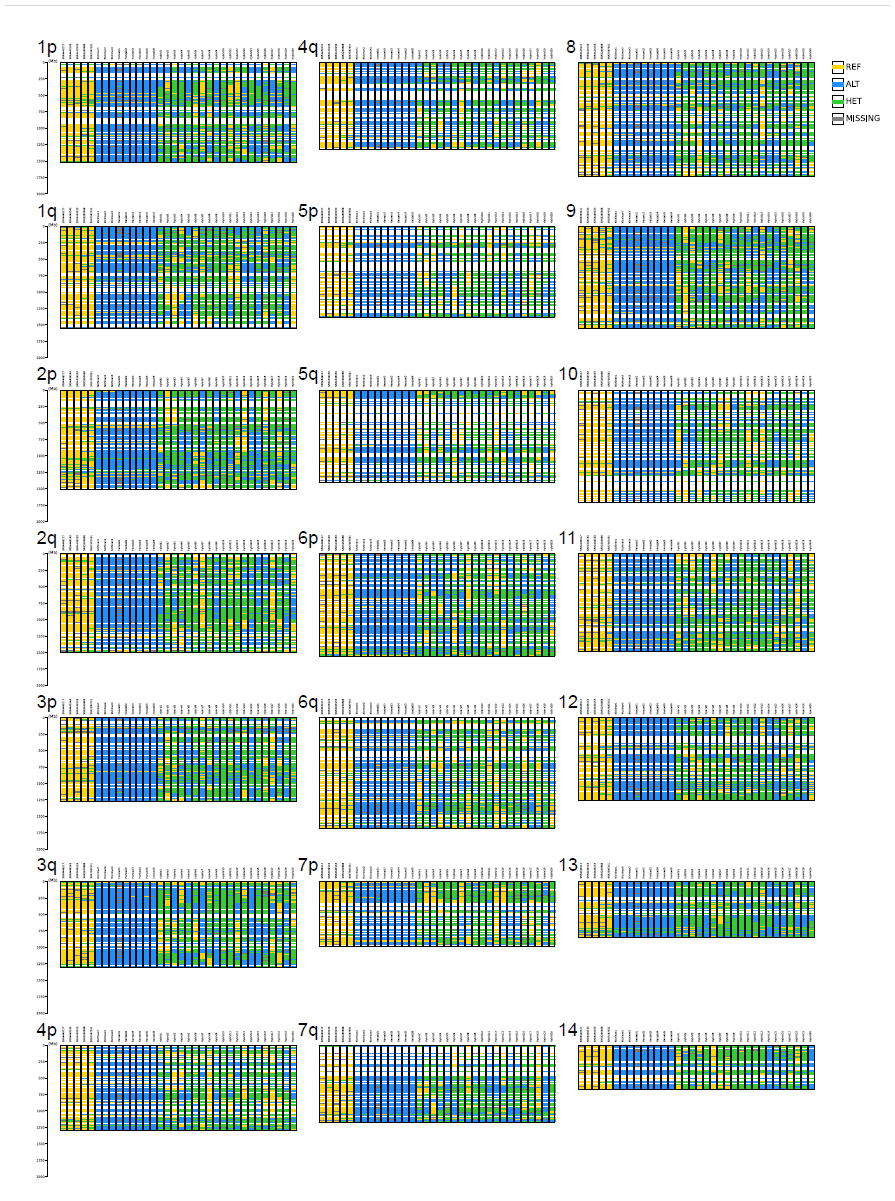

**Figure S2. Genome-wide polymorphism profile of the giant salamander based on pseudo-genome mapping.** An overview of the data presented in Table S11 is shown.

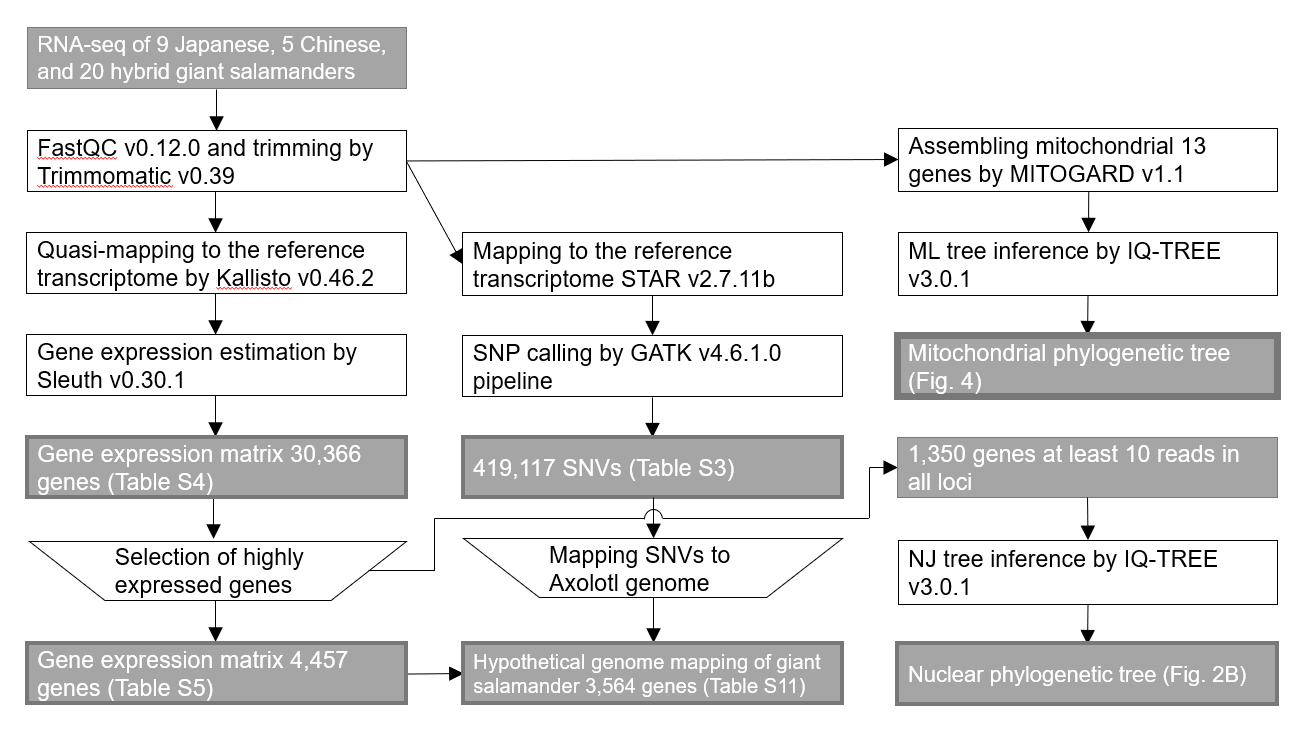

**Figure S3. Schematic overview of the analytical pipeline and software used in this study.**

**Supplementary Tables**

**Table S1. Total sequence reads used in this study**

**Table S2. Percentage of uniquely mapped reads for all 34 individuals**

**Table S3. A total of 419,177 SNP candidates identified among Japanese giant salamanders, Chinese giant salamanders, and hybrid individuals relative to the reference Chinese giant salamander genome.**

**Table S4. Expression profiles of all 30,366 genes in giant salamander skin**

**Table S5. Expression profiles of 4,457 highly expressed genes in giant salamander skin**

**Table S6. A total of 2,393 differentially expressed genes (DEGs) identified in hybrid skin relative to Japanese giant salamander skin, including 1,343 upregulated and 1,050 downregulated genes.**

**Table S7. Gene Ontology (GO) enrichment analysis of genes significantly upregulated in hybrid skin compared with Japanese giant salamander skin.**

**Table S8. Expression profiles of selected genes upregulated in hybrids**

**Table S9. Gene Ontology (GO) enrichment analysis of genes significantly downregulated in hybrid skin compared with Japanese giant salamander skin.**

**Table S10. Expression profiles of selected genes downregulated in hybrids**

**Table S11. Genome-wide polymorphism profile of the giant salamander based on pseudo-genome mapping**

**Table S12. Haplotypes of seven environmental adaptation-related genes and adjacent loci**

**Table S13. Chromosome sizes and IDs of the axolotl (*Ambystoma mexicanum*) genome**
